# Nutrient fluctuations alter effects of litter diversity of invasive species on native communities

**DOI:** 10.1101/2025.11.05.686688

**Authors:** Xue Wang, Mark van Kleunen, Wei-Long Zheng, Jing-Jing Han, Chun-Lan Wu, Shan Yang, Fei-Hai Yu

## Abstract

Invasive alien plants can indirectly suppress native plants by altering soil biota and nutrient cycling through their litter input. The diversity of litter resulting from co-invasion by multiple species may further modulate these impacts. Fluctuating resources are known to favour many invasive plants; however, it is unclear how nutrient fluctuations alter the effects of litter diversity of invasive plants on native communities. Therefore, we grew a native plant community in a control soil without litter, as well as in soils mixed with litter from one, two, three, and six invasive species under constant or pulsed nutrient supply. Invasive species litter altered soil microbial communities, and increased soil total nitrogen concentration and native community biomass. Under pulsed nutrient supply, native community biomass and soil total phosphorus concentration decreased with increasing litter diversity of the invasive species. However, these effects did not occur under constant nutrient supply. A structural equation model indicated that soil phosphorus and fungal community composition were key mediating factors driving the decrease in native community biomass with increasing litter diversity under pulsed nutrient supply. Our findings underscore that the impact of alien invaders on native communities depends on the diversity of the alien plant litter and nutrient fluctuations.

**Highlight:** Under pulsed nutrient supply, increasing invasive species litter diversity reduced native community biomass, mediated primarily by shifts in soil phosphorus and fungal community composition. These effects were absent under constant nutrient conditions.

## Introduction

Invasive plant species can alter the structure and functioning of ecosystems in their non-native ranges, causing tremendous ecological and economic damage (van Kleunen et al. 2015, Wang et al. 2017, Pyšek et al. 2020, Diagne et al. 2021). Invasive plants can directly suppress native plants by competing for resources and space (Wang et al. 2019, Lear et al. 2022, Chen et al. 2024) and can also indirectly inhibit native plants by producing allelochemicals and altering soil communities via their root exudates (Reinhart and Callaway 2006, Lu et al. 2018, Yu et al. 2022). Additionally, invasive plants may influence native plants through the introduction of their litter into the soil, which may modify the soil environment by introducing new pathogens (Massoni et al. 2021b, Benitez et al. 2022) and releasing nutrients and allelochemicals (Zhang et al. 2021, Semchenko et al. 2022). Previous studies have examined the impacts of litter inputs from invasive species on soil biota and nutrients and how these impacts feedback to influence invasion success (Inderjit et al. 2011, Tamura et al. 2017, Heras et al. 2020, Berta and Mott 2023). However, most of these studies have focused on single invasive plant species.

A common phenomenon is that native plant communities are being invaded by different numbers of alien plant species (McGrannachan and McGeoch 2019, Oschrin and Reynolds 2019, Sheppard 2019, Stotz et al. 2020). However, most theories of biological invasion are based on the invasion by a single invasive species (Callaway et al. 2004, Joshi and Vrieling 2005, Catford et al. 2009, Stotz et al. 2020, Guo et al. 2024). Although some studies have explored the influence of the co-invasion of two or three invasive plant species on native plant communities (Kuebbing et al. 2016, Mahla and Mlambo 2019, Oschrin and Reynolds 2019), the impact of invasive plant species diversity has been largely overlooked. Additionally, when a native community has been invaded by multiple alien species, their ecological threat persists long after active growth ends. Beyond direct competition, their mixed litter legacies—formed after death—create prolonged cascading effects on soil processes (Santonja et al. 2017, Bennett and Klironomos 2019). This knowledge gap is particularly urgent given real-world management challenges, where partial removal of invaders often leaves behind mixed litter. Such invasive plant legacies, no matter whether from monocultures or polycultures, could modify soil conditions with consequences for native plant recovery. Despite its implications for restoration, no study has yet tested how litter inputs from multiple invasive species jointly influence native plant communities.

Litter diversity of native plant species has been found to have positive, negative, or neutral effects on litter decomposition rates (Thomas 1968, Staaf 1980, Gartner and Cardon 2004, Gessner et al. 2010, Porre et al. 2020, Canessa et al. 2022). These mixed results might be a consequence of the different mechanisms that underlie litter diversity effects. A mixture of litter from a higher number of native plant species is likely to induce nutrient transfer among litter of different plant species and attract more diverse meso- and micro-fauna, which may accelerate litter decomposition (Handa et al. 2014, Bonanomi et al. 2017, Santonja et al. 2017, Otsing et al. 2018). In contrast, the transfer of toxic or recalcitrant compounds among litter of different plant species may negatively affect litter decomposition (McArthur et al. 1994, Freschet et al. 2012). As the litter decomposition rate is closely related to how fast nutrients and allelochemicals are released into the soil and the activity of meso- and micro-fauna, it will feedback to influence the growth of plant communities. As litter inputs from invasive species are commonly found to benefit the growth of invasive species more than that of native species (Mariotte et al. 2017, Zhang et al. 2019, Benitez et al. 2022), this effect may also be strengthened or weakened when litter diversity of invasive species is higher. However, it remains unknown whether litter diversity of invasive plant species influences native plant communities via its impact on soil biota and nutrients.

Nutrients are crucial for plant growth and are often released into the environment in pulses (Jentsch and White 2019, Gao et al. 2021). Nutrient fluctuation (e.g. a nutrient pulse) can benefit the growth of invasive species more than that of native species because invasive species frequently grow faster and have a greater ability to acquire resources (Parepa et al. 2013, Liu and van Kleunen 2017). Nutrient fluctuation may influence intraspecific and interspecific competition and consequently change species composition and productivity of plant communities (Edwards et al. 2013, Wang et al. 2015, Yuan et al. 2017, Zhang et al. 2022). Additionally, nutrient fluctuation may alter soil microbial abundance and activities, and thus, litter decomposition, which may further impact plant communities (Manzoni et al. 2010, Zhang et al. 2023). Therefore, nutrient fluctuation may interact with litter input to influence plant communities. However, to date, no study has tested the interactive effects of nutrient fluctuation and litter diversity of invasive plant species on native plant communities.

We grew a native plant community in a control soil without any litter application, as well as in soils mixed with litter from one, two, three, and six invasive species under constant or pulsed nutrient supply. We measured the productivity (biomass) of the native plant community and analysed the soil nutrient content and microbial composition. Specifically, we addressed the following questions: (1) Do litter inputs and litter diversity of invasive plants influence the productivity of the native plant community? (2) Does fluctuating nutrient availability modify the effects of litter inputs and litter diversity of invasive species on native plant community productivity? (3) Do soil microbes mediate the effects of litter diversity of invasive plant species and nutrient fluctuation?

## Materials and methods

### Litter of invasive species and seedlings of native species

In November 2021, we collected litter (withered shoots) from six invasive plant species, *Ageratum conyzoides* L. (Asteraceae), *Bidens frondosa* L. (Asteraceae), *Bidens pilosa* L. (Asteraceae), *Sesbania cannabina* (Retz.) Poir. (Fabaceae), *Solidago canadensis* L. (Asteraceae), and *Symphyotrichum subulatum* (Michx.) G. L. Nesom (Asteraceae), in Taizhou, Zhejiang Province, China. In Taizhou, these six invasive species are widespread and frequently co-occur in herbaceous plant communities in the mountainous areas, roadsides, agricultural fields, and urban environments (Wang et al. 2022b, Wu and Yu 2023). *Ageratum conyzoides*, *B. frondosa*, *B. pilosa*, *S. cannabina* and *S. subulatum* are annuals, whereas *S. canadensis* is a perennial. The litter was air-dried and ground into powder before use.

The native plant community consisted of two grass species (*Elymus dahuricus* Turcz. and *Lolium arvense* With.), two non-legume forbs (*Aster ageratoides* Turcz. and *Plantago asiatica* L.), and two legumes (*Astragalus sinicus* L. and *Medicago sativa* L.). We chose these native species because they frequently occur in herbaceous plant communities in the mountainous areas of Taizhou, and because their seeds are commercially available. Seeds of these native species were obtained from a local supplier in Taizhou. The seeds of the six native plants were sown in plastic pots (54 cm × 28 cm × 5 cm) filled with peat in a greenhouse at the Jiaojiang Campus of Taizhou University in Taizhou, Zhejiang Province, China, on 17 May 2022. Similarly sized seedlings of each species were selected on 11 June 2022 and used to construct the native plant community.

### Environmental design

The native plant community was constructed in 7.5 L pots filled with a control soil without any litter application or soils mixed with litter from one, two, three, or six invasive species under constant or pulsed nutrient supply. The control soil was a 1:1 (v/v) mixture of river sand and a field soil containing 0.09 ± 0.02 (mean ± SE) g kg^-1^ total nitrogen (N) and 0.03 ± 0.02 g kg^-1^ total phosphorus (P). The field soil was collected from a mountainous area where the invasive and the native plants are widely distributed. As the field soil was rich in clay, we mixed it with river sand to facilitate plant rooting and thus growth. For the soils with litter, 15 g of invasive species litter was mixed evenly into the soil in each pot, equivalent to a total concentration of 0.2% (w/w).

For the one-species litter treatment, the litter of each of the six invasive plant species was applied to the soil mixture in three pots of each nutrient treatment, resulting in 36 pots (6 invasive species × 2 nutrient treatments × 3 replicates). For the two-species litter treatment, litter mixtures from 12 different two-species combinations were randomly selected and applied to 12 pots for each nutrient treatment, resulting in 24 pots (12 litter mixtures × 2 nutrient treatments × 1 replicate). For the three-species litter treatment, litter mixtures from 12 different three-species combinations were randomly selected and applied to 12 pots for each nutrient treatment, resulting in 24 pots (12 litter mixtures × 2 nutrient treatments × 1 replicate). The two- and three-species litter mixtures were selected according to the criterion that the litter from each species should be used at the same frequency, similar to the design of many biodiversity experiments (Hector et al. 1999, Wang et al. 2022b). Due to the limited number of litter species, the litter mixture of all six invasive species was applied to 12 pots of each nutrient treatment for the six-species litter treatment, resulting in 24 pots (1 litter mixture × 2 nutrient treatments × 12 replicates). Twelve replicates (pots) of each nutrient treatment were used as a control (no litter application), resulting in 24 pots. So, there were 132 pots in total. An equal amount of litter (15 g) was added to the soil mixture in each pot. Thus, 15, 7.5, 5 and 2.5 g litter from each species was mixed into the soil for the one-, two-, three-, and six-species litter treatments, respectively.

A native plant community was constructed in each pot on 11 June 2022 by transplanting two seedlings of each of the six native plant species. Seedlings that died were immediately replaced within the first 2 weeks after transplantation. Three weeks after transplantation i.e., on 4 July 2022, we started the nutrient treatments using a 4× strength Hoagland solution (945 mg L^-1^ Ca(NO_3_)_2_·4H_2_O, 607 mg L^-1^ K_2_SO_4_, 115 mg L^-1^ NH_4_H_2_PO_4_, 493 mg L^-1^ MgSO_4_, 20 mg L^-1^ Na_2_-EDTA, 2.86 mg L^-1^ FeSO4, 4.5 mg L^-1^H_3_BO_3_, 2.13 mg L^-1^ MnSO_4_, 0.05 mg L^-1^ CuSO_4_, 0.22 mg L^-1^ ZnSO_4_, 0.02 mg L^-1^ (NH_4_)_2_SO_4_). During the experiment, 200 mL of Hoagland solution was added to the pots. For the constant nutrient treatment, 20 mL of nutrient solution was supplied to each pot weekly for 10 weeks. For the pulsed nutrient treatment, we supplied 8 mL of the nutrient solution to each pot per week for the first 3 weeks, then 28 mL per week in the following 4 weeks, and again 8 mL per week for the next 3 weeks. We added extra water to the nutrient solution in each treatment to ensure that each pot received 100 mL of water per nutrient application and avoid differences in the water supply between the two nutrient treatments.

### Plant harvest and soil sampling

At the end of the experiment, we harvested the shoots and roots of each native plant species from each pot. Shoots and cleaned roots were oven-dried at 65 °C for 72 hours and then weighed. Two mixed soil samples were collected for each pot. One soil sample was stored at −20 °C and used for soil microbial community composition analysis; the other soil sample was air-dried and used for soil pH, total N, and total P analyses.

### Measurements of soil physicochemical properties

Soil pH was determined using a 1:2.5 soil-to-water ratio (w/v). Total N and P concentrations were measured using an Auto Analyzer 3 (Bran and Luebbe, Norderstedt, Germany) following acid digestion with a 10:1 (v/v) mixture of sulfuric and perchloric acids.

### Soil microbial community analysis

Soil DNA extraction, amplification, library preparation, sequencing, and bioinformatics analyses were conducted by Majorbio Co., Ltd. (Shanghai, China). The libraries were constructed using PCR with the TransStart Fastpfu DNA Polymerase kit (AP221-02, TransGen, Beijing, China) in a PCR cycler (GeneAmp 9700, ABI, Carlsbad, California, USA). The primers 338F-ACTCCTACGGGAGGCAGCAG and 806R-GGACTACHVGGGTWTCTAAT (Liu et al. 2016) were used to amplify the V4 region of the bacterial 16S rDNA, whereas ITS1F-CTTGGTCATTTAGAGGAAGTAA and ITS2R-GCTGCGTTCTTCATCGATGC (Gardes and Bruns 1993) were used to amplify the fungal ITS rDNA region. After amplification and recovery, the PCR products were mixed in the proportions required for sequencing. Libraries were generated using a TruSeq DNA Sample Prep Kit and sequenced on an Illumina NovaSeq 6000 platform (Illumina, San Diego, California, USA). After sample splitting of the paired-end reads obtained using NovaSeq sequencing, the double-end reads were first quality-controlled and filtered according to sequencing quality, spliced according to the overlap relationship, and optimised data were obtained. Finally, the optimised data were processed using sequence denoising methods (e.g. DADA2/Deblur) to obtain amplicon sequence variant (ASV) sequences and abundance information. Classify-sklearn (Naive Bayes), classify-consensus-vsearch (Vsearch), classify-consensus-blast (Multi_Blast), and RDP classifier Bayes algorithms were used to analyse the ASV sequences and obtain the species classification information for each ASV. The BLAST databases were Silva (for bacteria and archaea, Release138, http://www.arb-silva.de) and Unite (for fungi, Release 8.0 https://unite.ut.ee/). The sequence numbers of each sample were rarified to 41,762 reads to minimise the effects of sequencing depth on community diversity (alpha and beta diversity). Putative fungal functional groups (e.g. pathogenic fungi) were identified using the FUNGuild database (Nguyen et al. 2016).

### Statistical analysis

All analyses were performed using R version 4.3.3 (R Core Team 2024). Linear mixed models were used to test the effects of litter application (absent vs present, where the latter is averaged over one, two, three, and six invasive species), nutrient fluctuation (constant vs pulsed), and their interaction on the biomass (total, shoot, and root) of the native plant community; soil pH, total N, and total P concentrations; the relative abundance of the top 10 bacterial and fungal communities at the phylum and class levels; and the relative abundance of the top 10 plant pathogenic fungi at the family level. Litter species composition was included as random factors in these models. Subsequently, the no-litter treatment was omitted from the analysis that focused exclusively on effects of litter diversity. Thus, linear mixed models were used to test the effects of litter diversity (litter from one, two, three, and six invasive species), nutrient fluctuation, and their interactive effects on the variables. Litter species composition was treated as a random factor in these models. Data on the biomass of the native plant community and the relative abundance of the top 10 pathogenic fungi at the family level were log-transformed to improve the normality and homoscedasticity of the residuals. Duncan’s test was used for multiple comparisons.

To assess potential effects of litter application or litter diversity, nutrient fluctuation, and their interactions on the final composition of the native community, we assessed variation in plant species composition, based on biomass proportions, using principal component analysis (PCA). For each PCA (Fig. S1), we extracted the PC1 and PC2 values, which together explained more than 58% of the variation in community composition, and included them as response variables in separate linear mixed models, as described above for community biomass.

Bray–Curtis distances of microbial community composition were computed using the vegan package based on ASV tables and visualised using nonmetric multidimensional scaling (NMDS) plots. Permutational multivariate analysis of variance (PERMANOVA) was performed using the ADONIS function based on the Bray–Curtis dissimilarity matrix to determine whether litter application and litter diversity under both constant and pulsed nutrient treatments significantly influenced microbial community composition.

Structural equation modelling (SEM) was applied using the lavaan package (Rosseel 2012) in R to explore how increasing litter diversity under pulsed nutrients affected the total and shoot biomass of the native community via effects on soil P, soil pH, and composition of soil fungal communities. This was only done for the pulsed treatment, because litter diversity only had a significant effect in that treatment. A direct effect of litter diversity was not included in the SEM, because if there is a direct effect, it might not be a true direct effect, but an effect mediated by unknown factors not included in the model. We assume that litter diversity might affect total P, pH, and composition of fungal communities in the soil, and subsequently influence the total and shoot biomass. The composition of soil fungal communities was indicated by the first and second nonmetric multidimensional scaling axes in models. The fit of the model to the data was determined using the χ^2^ test, goodness-of-fit index (GFI) and root-mean-square error of approximation (RMSEA).

A significant effect of litter diversity may stem from two mechanisms: (1) the increased probability of including litter from an invasive plant species with strong competitive dominance, or (2) synergistic interactions between litter from different invasive species, which could either depend on specific species combinations or represent generalized interaction effects. To test these hypotheses, we applied the hierarchical diversity interaction modeling framework of Kirwan et al. (2009) to distinguish between identity effects (individual species contributions) and interaction effects (species interactions) driving litter diversity impacts on total and aboveground biomass under pulsed nutrients. Five hierarchical models were evaluated for both biomass metrics, differing in their assumptions about the contributions of individual invasive species’ litter effects and their interactions to native community productivity. The best fitting model was identified using likelihood ratio tests (see Method S1 for details).

## Results

### Native plant community productivity

The presence of litter (i.e. averaged over applications of litter from one, two, three, and six invasive plant species) significantly increased total biomass (Fig. 1a, upper right corner) and root biomass (Fig. 1c, upper right corner) and marginally significantly increased shoot biomass (Fig. 1b, upper right corner) of the native community. The effects of litter diversity (i.e. litter from one, two, three, and six invasive plant species) on total and shoot biomass were significantly influenced by nutrient fluctuation. Specifically, increasing litter diversity led to a significant decrease in total and shoot biomass under pulsed nutrient supply, whereas no changes were observed under constant nutrient supply (Fig. 1a, b, main graphs). Root biomass (Fig. 1c, main graph) increased significantly with increasing litter diversity. However, neither nutrient fluctuation nor its interaction with litter diversity significantly affected root biomass in the native plant community (Fig. 1c, main graph).

**Figure 1.**
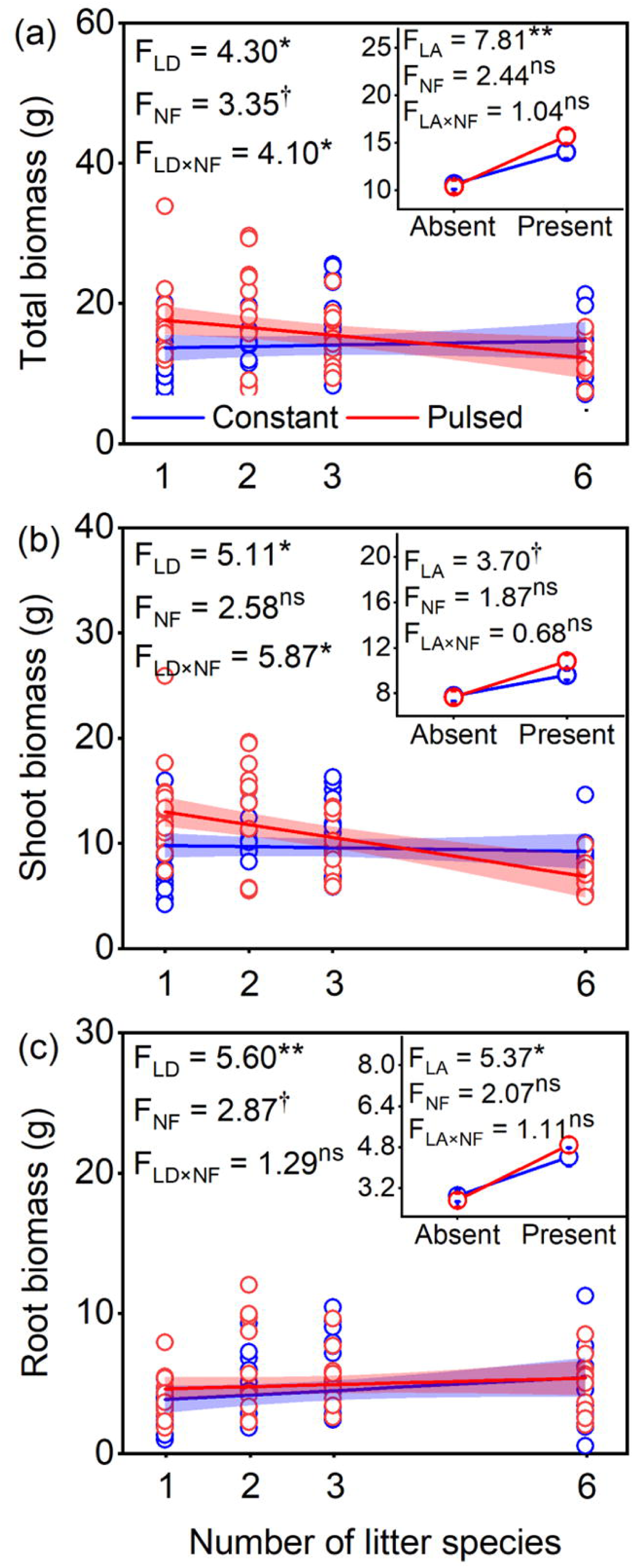
The response of the productivity of the native plant community to the interactive effects of litter application or diversity and nutrient fluctuation. (a) Total biomass. (b) Shoot biomass. (c) Root biomass. In the main graph, solid lines are the fitted relationships for the effects of litter diversity (litter from 1, 2, 3 and 6 invasive plant species) and nutrient fluctuation (constant vs pulsed), and the coloured polygons delimit the 95% confidence intervals. Insets at the upper right corner of each graph show the effects of litter application (absent vs present, averaged over litter from 1, 2, 3 and 6 invasive species) and nutrient fluctuation. F-values of linear mixed models are given. The asterisks (*) and (**) indicate significant differences at *P* < 0.05 and *P* < 0.01, respectively. The daggers (†) indicate marginally significant differences at *P* < 0.1, and the letters ‘ns’ indicate non-significant values. Model terms ‘LD’, ‘LA’ and ‘NF’ indicate litter diversity, litter application and nutrient fluctuation, respectively.

Hierarchical diversity interaction modelling revealed that the model incorporating separate pairwise interactions provided the best fit for total biomass and shoot biomass under pulsed nutrients, indicating that the effect of litter diversity on these parameters was mainly driven by specific pairwise interactions (Table 1).

**Table 1.** Contributions of invasive plant species (IPS) identity and interactions to the litter-diversity effect on total biomass and shoot biomass under pulsed nutrients. Comparison of the hierarchical diversity interaction models (Methods S1) were based on log-likelihood ratio tests to assess the contributions of individual IPS and IPS-interactions to the litter diversity effect on biomass. The statistics (AIC, logLik, χ^2^ and *P*) of the best fitting models (i.e. with the lowest AIC) are highlighted in bold.

| Model (reference model number) | Total biomass |  |  |  |  | Shoot biomass |  |  |  |
| --- | --- | --- | --- | --- | --- | --- | --- | --- | --- |
| | Df | AIC | logLik | $\chi^2$ | $P$ | AIC | logLik | $\chi^2$ | $P$ |
| 1. Null | - | 41.71 | -16.86 | - | - | 36.09 | -14.04 | - | - |
| 2. Invasive plant species (IPS) identity (1) | 5 | 51.6 | -16.8 | 0.11 | 0.999 | 42.65 | -12.33 | 3.43 | 0.633 |
| 3. Separate-pairwise IPS interactions (4) | 14 | **18.52** | **14.74** | **63.28** | **<0.001** | **7.74** | **20.13** | **65.42** | **<0.001** |
| 4. Average IPS interaction (2) | 1 | 53.8 | -16.9 | 0.19 | 0.659 | 45.16 | -12.58 | 0.51 | 0.473 |
| 5. Additive IPS-specific interaction contributions (3) | 9 | 51.51 | -10.78 | 50.99 | <0.001 | 42.18 | -6.09 | 52.44 | <0.001 |

### Native plant community composition

To explicitly test whether the different responses of the native species affected community composition, we used PCA analysis. In the native community exposed to litter application and nutrient fluctuation, the first two axes of the PCA explained 58.2% of the variance in species composition (Fig. S1a). In the native community exposed to increasing litter diversity and nutrient fluctuation, the first two axes of the PCA explained 62.9% of the variance (Fig. S1b). Litter application had no significant effects on values of PC1 and PC2 (Fig. S2, upper right corner). Nutrient fluctuation significant decreased the value of PC2 but did not significantly affect the value of PC1 (Fig. S2, upper right corner and main graph). Increasing litter diversity significantly increased the PC1 value (Fig. S2a, main graph) but had no effects on the PC2 value (Fig. S2b, main graph).

### Soil nutrients and pH

Total soil N was significantly lower under pulsed than under constant nutrient supply, and litter application significantly increased total soil N (Fig. 2a, upper right corner). Litter diversity did not affect total soil N (Fig. 2a, main graph). On the other hand, litter application led to a significant decrease in total soil P, but only under pulsed nutrient supply, not under constant nutrient supply (Fig. 2b, upper right corner). Total soil P significantly decreased with increasing litter diversity of the invasive plant species under pulsed nutrient supply but not under the constant nutrient supply, as indicated by a significant two-way interaction between litter diversity and nutrient fluctuation (Fig. 2b, main graph). Litter application significantly increased soil pH under constant nutrient supply but not under pulsed nutrient supply (Fig. 2c, upper right corner). Soil pH did not vary with litter diversity, but it was significantly lower under pulsed nutrient supply than under constant nutrient supply (Fig. 2c, main graph).

**Figure 2.**
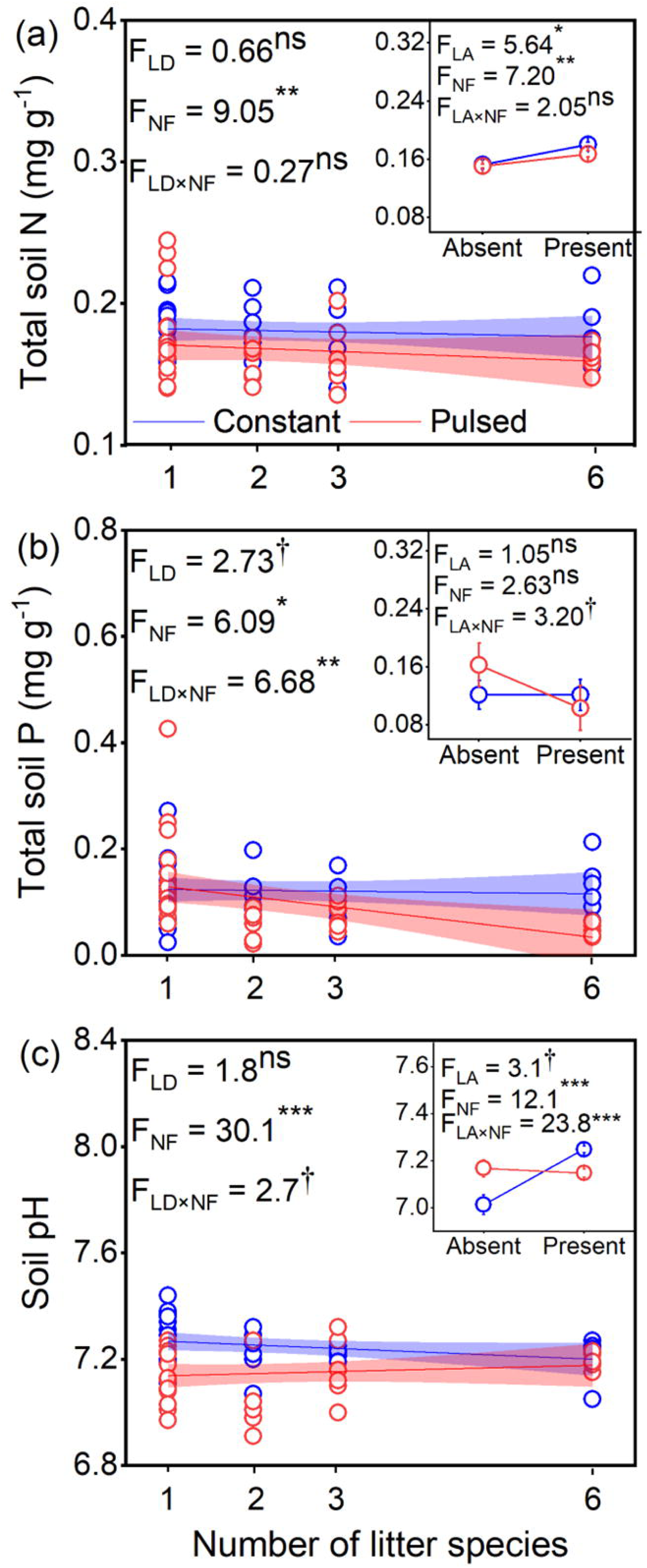
The response of soil nutrients to the interactive effects of litter application or diversity and nutrient fluctuation. (a) The concentration of total soil N. (b) The concentration of total soil P. (c) Soil pH. In the main graph, solid lines are the fitted relationships for the effects of litter diversity (litter from 1, 2, 3 and 6 invasive plant species) and nutrient fluctuation (constant vs pulsed), and the coloured polygons delimit the 95% confidence intervals. Insets at the upper right corner of each graph show the effects of litter application (absent vs present, averaged over litter from 1, 2, 3 and 6 invasive species) and nutrient fluctuation. F-values of linear mixed models are given. The asterisks (*), (**) and (***) indicate significant differences at *P* < 0.05, *P* < 0.01 and *P* < 0.001, respectively. The daggers (†) indicate marginally significant differences at *P* < 0.1, and the letters ‘ns’ indicate non-significant values. Model terms ‘LD’, ‘LA’ and ‘NF’ indicate litter diversity, litter application and nutrient fluctuation, respectively.

### Soil microbial community

Litter application, litter diversity, and nutrient fluctuation significantly altered the composition of the bacterial (Fig. 3a, c) and fungal (Fig. 3b, d) communities. Moreover, the composition of the bacterial community was also affected by the interaction between litter application and nutrient fluctuation (Fig. 3a). The relative abundances of the dominant bacterial (at the phylum and class levels) and fungal (at the phylum, class, and major plant-pathogenic family levels) taxa are detailed in Result S1.

**Figure 3.**
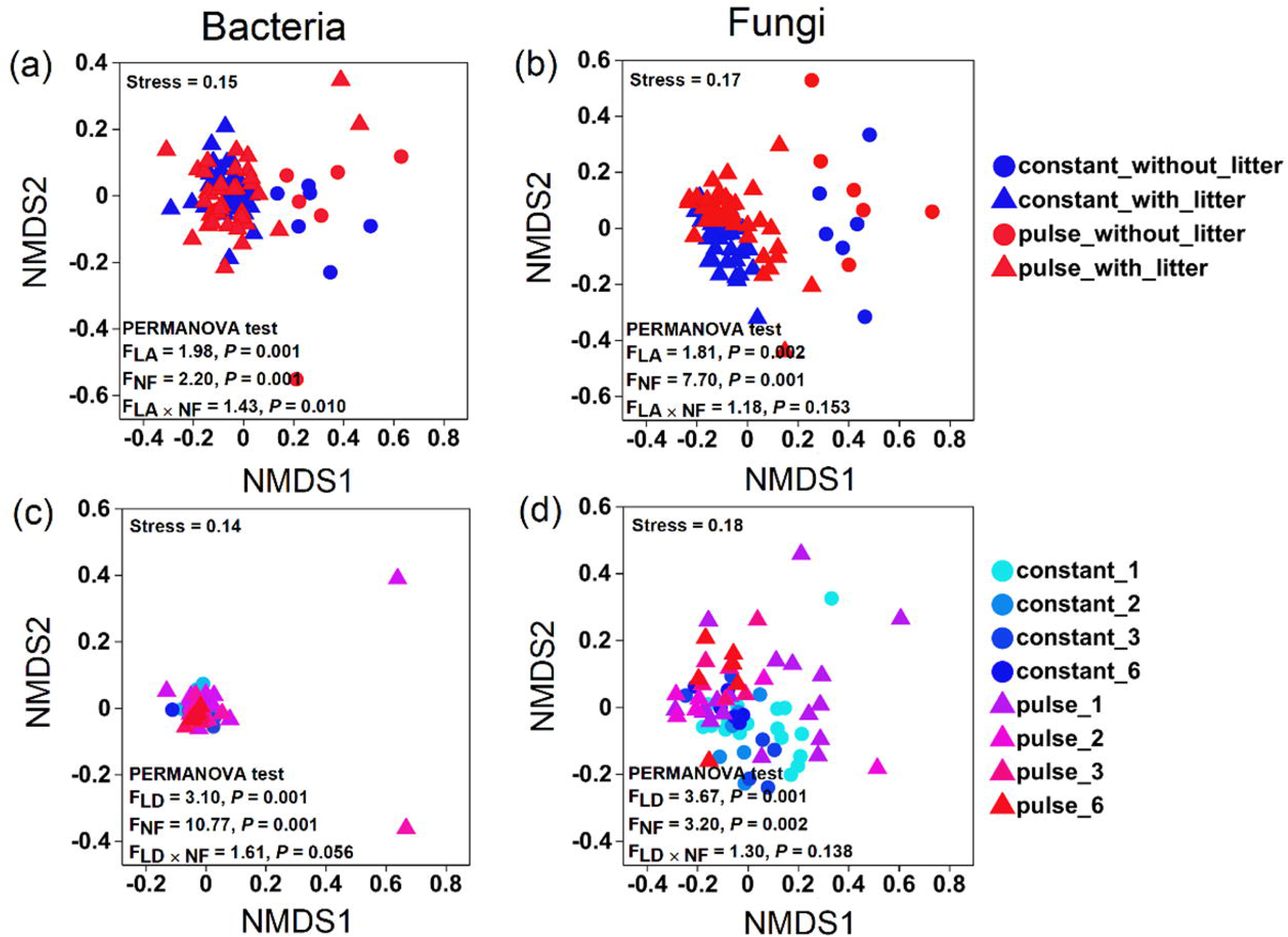
The soil microbial composition. (a) The bacterial community composition under the effects of litter application (absent vs present, averaged over litter from 1, 2, 3 and 6 invasive species) and nutrient fluctuation (constant vs pulsed). (b) The fungal community composition under the effects of litter application and nutrient fluctuation. (c) The bacterial community composition in the treatments with different litter diversity levels (litter from 1, 2, 3 and 6 invasive species) and nutrient fluctuation. (d) The fungal community composition under different litter diversity levels and nutrient fluctuation. The compositions of the microbial communities are illustrated using nonmetric multidimensional scaling (NMDS) plots based on Bray–Curtis distances. Model terms ‘LD’, ‘LA’ and ‘NF’ indicate litter diversity, litter application and nutrient fluctuation, respectively.

### Effects of litter diversity on community total and shoot biomass through different mechanistic pathways under pulsed nutrient supply

The above analyses only found effects of litter diversity on various variables in the pulsed nutrient treatment. Therefore, to discriminate the indirect effects of litter diversity, soil P, pH, and fungal community composition on total and shoot biomass, we used a structural equation model for the pulsed nutrient condition only. The model fit was generally good (*P* > 0.05, RMSEA close to 0, and GFI close to 1). Increasing litter diversity negatively influenced soil P, and soil P subsequently had marginally positive effects on total biomass (Fig. 4a) and significantly positive effects on shoot biomass (Fig.4b). Litter diversity tended to negatively influence the NMDS1 score but had positive effects on the NMDS2 score of the fungal community composition (Fig. 4a, b). Both NMDS1 and NMDS2 scores had negative effects on total and shoot biomass (Fig. 4a, b).

**Figure 4.**
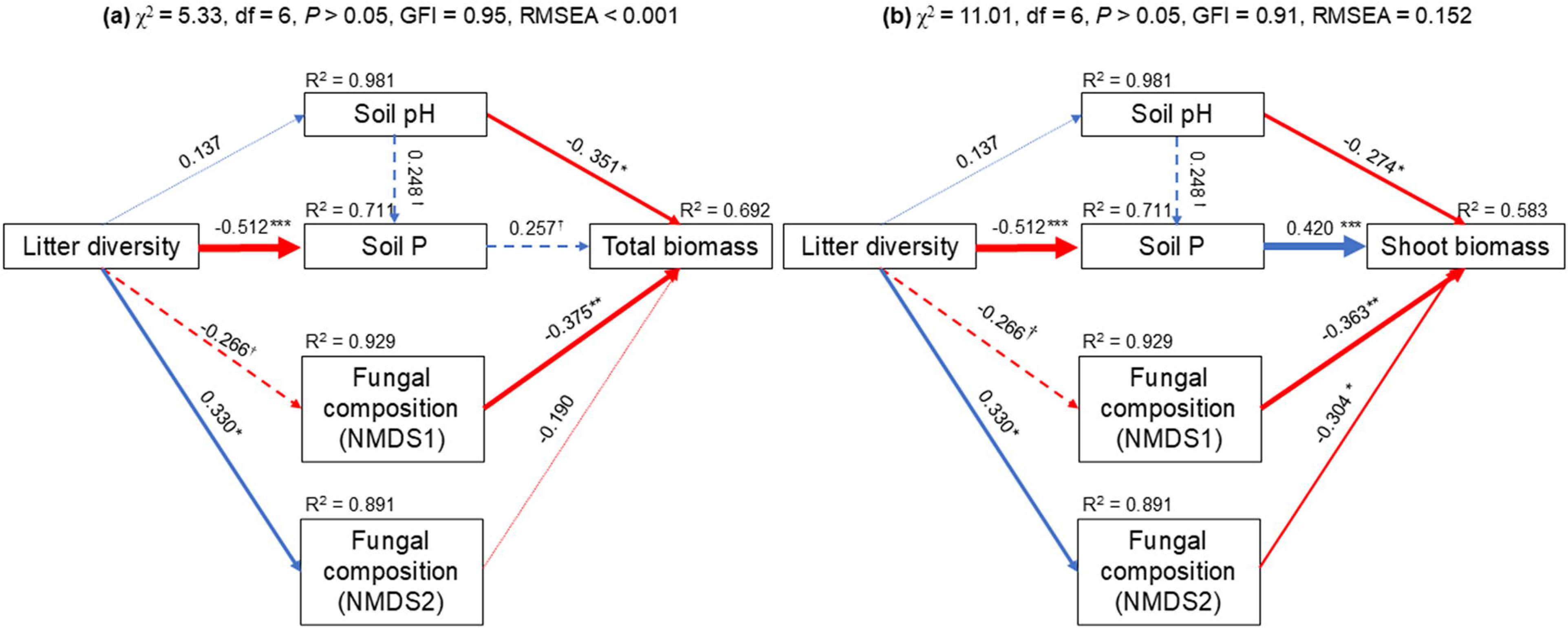
Structural equation modelling the indirect pathways of the litter diversity effect under pulsed nutrients on (a) total and (b) shoot biomass. Solid blue arrows indicate significant positive relationships, and solid red arrows indicate significant negative relationships (*P* < 0.05). Dashed arrows indicate marginally significant relationships (*P* < 0.1), and dotted arrows indicate non-significant relationships (*P* > 0.1). Numbers adjacent to the arrows represent standardised path coefficients. *R*^2^ values close to the variables indicate the variance explained by the model. ^†^0.1< *P* < 0.05, *0.01 < *P* < 0.05, **0.001 < *P* < 0.01, *** *P* < 0.001.

## Discussion

The impacts of invasive plants on native plant communities have been found to increase with the number of invasive species (Wang et al. 2022b). This is most likely due to increased competition (Wang et al. 2022b) and changes in soil communities (Wang et al. 2022a). However, to the best of our knowledge, no study had yet evaluated how litter diversity of invasive plant species influences native plant communities, i.e., legacy effects of mixed litter of invasive plant species after their death, and whether such a diversity effect is modified by nutrient fluctuations. Our study showed that while the productivity of native plant communities increased following the addition of invasive plant litter, this positive effect was reversed as litter diversity increased—but only under pulsed nutrient supply.

Specifically, we observed a 31.7% increase in total biomass following the application of invasive plant litter, which was mostly accounted for by a 70.6% increase in shoot biomass (Fig. 1). These results indicate that invasive plant litter does not exert negative effects due to, for example, the influence of pathogens or toxic effects of allelopathic chemicals (Bennett and Klironomos 2019, Massoni et al. 2021a), or that any negative effects were outweighed by positive effects due to the nutrients released from the litter (Meisner et al. 2012). Indeed, in our study, soil N increased significantly with litter application (Fig. 2a), thereby providing a nutrient-rich environment for plant growth.

Litter also supplies energy and nutrients to the soil biota, which can result in pronounced changes in soil microbial composition, including an increase in decomposers (He et al. 2023). The latter speed up nutrient cycling and thereby increase the abundance of nutrients available for plant growth (Veen et al. 2019). Our study revealed significant effects of litter application on soil microbial composition (Figs. 3, S3, S4). Specially, litter application significantly increased the relative abundances of fungal classes and phyla with many decomposers (Figs. S3, S4).

Although the presence of litter increased the biomass of the native community, the total biomass and shoot biomass were significantly reduced by 10.43% and 27.31%, respectively, when litter diversity increased from one to six invasive species (Fig. 1a, b, main graph). This was partly accounted for by different responses of the native species, as indicated by the strong effect of litter diversity on plant community composition (Figs. S1, S2). The litter diversity effects could be due to dilution of strong litter effects of certain invasive species or due to interactions between the litter produced by different invasive species. The hierarchical diversity interaction modelling showed that models that accounted only for the identities of the invasive plant species did not perform better than the null models. Therefore, the negative effect of litter diversity on native community biomass was most likely not due to dilution effects. The best models were the ones with separate pairwise interactions (Table 1), indicating that specific interactions between litter from different invasive plant species contributed to the effect of litter diversity on the productivity of the native community. These findings suggest that the combined litter inputs from multiple invasive species interact in ways that produce new ecological outcomes that are not predictable based on the effect of each species alone. This aligns with the growing recognition that interactions between multiple invaders often lead to complex, unpredictable outcomes (Kuebbing et al. 2013, Vujanović et al. 2022, Oduor et al. 2024). For example, in mixed litter environments, litter from one species might enhance the decomposition of the litter of another species, leading to changes in nutrient availability that affect plant growth (Gartner and Cardon 2004)

Nutrient fluctuation had a limited effect on the productivity of the native plant community and its effect depended on litter diversity. However, we found that nutrient fluctuation affected the composition of the native plant community (Fig. S2b). Nutrient fluctuation has been shown to strongly influence soil microbial composition (Carrero-Colon et al. 2006, Buscardo et al. 2018), and this could affect litter decomposition and nutrient release. In our study, pulsed nutrient conditions resulted in a significant decrease in the relative abundance of the dominant bacteria phylum Actinobacteriota (−13.0%) and the dominant fungus phylum Ascomycota (−10.1%) (Tables S3 and S4; Fig. S4), both of which include many decomposers. The reduction in these soil decomposers could lead to a decrease in nutrient release, which was supported by the significantly lower levels of total soil N at all litter diversity levels and a lower total soil P at high litter diversity under pulsed nutrient supply (Fig. 2a, b). Alternatively, a nutrient pulse may cause an imbalance in the availability of N and P, which can disrupt microbial communities, particularly decomposers that require both elements for efficient decomposition and nutrient release (Li et al. 2023, Qiu et al. 2023). A shift in nutrient availability can trigger microbial feedback loops, where the reduction in certain decomposer groups may limit nutrient mineralization, further restricting nutrient availability for plants (Encinas-Valero et al. 2024). Therefore, our findings indicate that nutrient fluctuation can negatively affect the performance of native plant communities, particularly at high litter diversity, by decreasing the relative abundance of key saprotrophic bacteria and fungi.

Our data revealed significant interactions between litter diversity and nutrient fluctuation in shaping the productivity of the native plant community. Total and shoot biomass showed no variation with increasing litter diversity under constant nutrient conditions. However, both total and shoot biomass decreased with increasing litter diversity under pulsed nutrient conditions (Fig. 1a, b). Our finding that total soil P decreased with increasing litter diversity under pulsed nutrient conditions (Figs. 2b and 4) suggests that the decrease in plant biomass under those conditions is due to nutrient limitation. Different taxa or functional groups within the soil microbial community have varying effects on nutrient cycling, and certain fungal groups have a higher ability to mobilise P. Changes in the abundance of these groups can greatly influence P mineralisation and assimilation (Smith and Read 2008). For example, Ma et al. (2023) suggested that Mortierellomycota may play a key role in P mineralisation and assimilation. However, the abundance of this phylum did not change with increasing litter diversity in our study (Table S4). This suggests that other microbial groups may be important for regulating P mineralisation under pulsed nutrient conditions.

Although the relative abundance of the dominant fungus phylum Ascomycota (∼80%) did not vary with increasing litter diversity (Table S4; Fig. S4d), the relative abundance of the major class Sordariomycetes increased significantly with increasing litter diversity under pulsed nutrient conditions (Table S2; Fig. S3d). The latter should benefit plant growth because it includes many saprotrophes. However, there is enormous phylogenetic and physiological diversity within each class, and it is unlikely that all fungi within the same class will share common ecological characteristics (Taylor et al. 2014, Lücking et al. 2021).

The results on the abundance of plant pathogenic fungi showed that the three most dominant pathogenic fungus families, Plectosphaerellaceae, Nectriaceae, and Hypocreales fam Incertae sedis, accounted for 59% of the total sequences. They all belonged to the class Sordariomycetes and the phylum Ascomycota, the sum of which tended to decrease under constant nutrient conditions but increased under pulsed nutrient conditions with increasing litter diversity (Fig. S5b). In addition, the bacterial class Bacillus, which includes many beneficial bacteria, marginally decreased under the pulsed nutrient conditions (Fig. S3b). This reduction in Bacillus could weaken pathogen suppression, allowing more plant pathogenic fungi to thrive and potentially reducing native plant biomass (Dimkić et al. 2022, Yang et al. 2024). These factors might contribute to a reduction in the native plant community by increasing litter diversity under pulsed nutrient conditions. Our results suggest that the composition of the fungal community strongly influences the effects of litter diversity on the native community biomass under pulsed nutrient conditions (Fig. 4).

We conclude that litter from multiple invasive plant species has significant interaction effects on the productivity of the native plant community. Litter application significantly increased the total biomass of the native plant community, but this positive effect significantly decreased with increasing litter diversity under pulsed nutrient conditions. P limitation and changes in fungal community composition (e.g. saprotrophic fungi and potentially pathogenic fungi) might contribute to biomass reduction with increasing litter diversity under pulsed nutrient conditions. Therefore, our findings suggest that nutrient fluctuations can modify the effects of litter diversity of invasive plant species on native plant communities.

## Aacknowledgements

This study was supported by the National Key RandD program (2023YFE0124900), National Natural Science Foundation of China (32401307) and the Taizhou Scientific and Technological Project (23nya19).

## Author contributions

Xue Wang and Fei-Hai Yu designed the study. Jing-Jing Han and Chun-Lan Wu, performed the experiment. Xue Wang, Wei-Long Zheng and Shan Yang analyzed the data. Xue Wang wrote the first draft. Mark van Kleunen and Fei-Hai Yu helped to improve the final draft. All authors read and approved the final manuscript.

## Notes

### Competing Interest Statement

The authors have declared no competing interest.

